# Object-based analyses in FIJI/ImageJ to measure local RNA translation sites in neurites in response to Aβ1-42 oligomers

**DOI:** 10.1101/2020.01.27.921494

**Authors:** María Gamarra, Maite Blanco-Urrejola, Andreia F.R. Batista, Josune Imaz, Jimena Baleriola

## Abstract

Subcellular protein delivery is especially important in signal transduction and cell behavior, and is typically achieved by localization signals within the protein. However, protein delivery can also rely on localization of mRNAs that are translated at target sites. Although once considered heretical, RNA localization has proven to be highly conserved in eukaryotes.

RNA localization and localized translation are especially relevant in polarized cells like neurons where neurites extend dozens to hundreds of centimeters away from the soma. Local translation confers dendrites and axons the capacity to respond to their environment in an acute manner without fully relying on somatic signals. The relevance of local protein synthesis in neuron development, maintenance and disease has not been fully acknowledged until recent years, partly due to the limited amount of locally produced proteins. For instance, in hippocampal neurons levels of newly-synthesized somatic proteins can be more than 20-30 times greater than translation levels of neuritic proteins. Thus local translation events can be easily overlooked under the microscope.

Here we describe an object-based analysis used to visualize and quantify local RNA translation sites in neurites. Newly-synthesized proteins are tagged with puromycin and endogenous RNAs labelled with SYTO. After imaging, signals corresponding to neuritic RNAs and proteins are filtered with a Laplacian operator to enhance the edges. Resulting pixels are converted into objects and selected by automatic masking followed by signal smoothing. Objects corresponding to RNA or protein and colocalized objects (RNA and protein) are quantified along individual neurites. Colocalization between RNA and protein in neurites correspond to newly-synthesized proteins arising from localized RNAs and represent localized translation sites. To test the validity of our analyses we have compared control neurons to Aβ1-42-treated neurons. Aβ is involved in the pathology of Alzheimer’s disease and was previously reported to induce local translation in axons and dendrites which in turn contributes to the disease. We have observed that Aβ increases the synthesis of neuritic proteins as well as the fraction of translating RNAs in distal sites of the neurite, suggesting an induction of local protein synthesis. Our results thus confirm previous reports and validate our quantification method.

## Introduction

Among all cell types, neurons are the most morphologically complex. The nucleus is contained in a cell body or soma, from where several neurites emerge. Neuronal dendrites measure around ten millimeters and axons can reach one meter of length in vertebrates (Bannister and Larkman, 1995b). This extremely polarized morphology reflects the also polarized function of neurons. Whereas dendrites receive signals, the cell body processes them and axons are responsible for transmitting information to adjacent neurons. To maintain a proper function, each neuronal compartment needs to react temporally and spatially in an acute manner in order to rapidly adapt to changes in the environment. These implies that compartmentalized signalling events are required and therefore neuronal proteins must be asymmetrically distributed.

The origin of neuritic proteins (both dendritic and axonal) has been discussed for years. It was classically thought that proteins that support dendritic and axonal functions are synthesized in the soma and then transported to the target compartment at peripheral sites of the neuron. However, in the 19^th^ century, the possibility of neurites, especially axons, producing their own proteins locally was already hypothesized (review in (Bolton, 1901)). This unconventional view of protein distribution to different neuronal compartments has been finally accepted by the scientific community. In order to synthesize proteins locally, messenger RNAs (mRNAs) and components of translational machinery must be transported to neurites. mRNAs are localized to dendrites and axons as part of ribonucleoprotein (RNPs) complexes in a translationally repressed state. Exogenous stimulus sensed by neurites influence the local translation machinery and mRNAs are released from RNPs complexes. Once associated to localized ribosomes, mRNAs are translated and proteins are synthesized independently from the soma and thus the endoplasmic reticulum (ER) (Jung et al., 2012).

The requirement of local intra-dendritic translation for nervous system plasticity has been extensively studied. Local translation in axons is involved in growth cone behaviour, axonal pathfinding and maintenance, as well as in retrograde signalling (reviewed in (Jung et al., 2014; Holt et al., 2019)). More recently, it has been reported that adult axons are also able to respond to pathological insults by changing their local translatome. In particular, after a nerve injury, mRNAs are locally translated and newly synthesized proteins contribute to axonal regeneration (Terenzio et al., 2018). Similarly, in the central nervous system (CNS) intra-axonal protein synthesis induced by Aβ_1-42_ oligomers, whose accumulation is central to Alzheimer’s disease (AD), contributes to neurodegeneration (Baleriola et al., 2014; Walker et al., 2018). Interestingly some authors have linked intra-dendritic translation and Tau mislocalization and hyperphosphorilation (Kobayashi et al.,2017; Li and Gotz, 2017). Thus, dysregulation of local protein synthesis might play a more relevant role in nervous system dysfunction than previously acknowledged.

AD is characterized by synaptic dysfunction during early stages (Palop and Mucke,2010). Understanding dynamic early changes in the local proteome is in our view crucial to understand basic pathological mechanisms underlying AD and likely other neurological diseases. An accurate quantification of local translation events, which is the aim of this study, might therefore give important clues to the extent to which changes in the local translatome contribute to the disease. Currently the most frequently used techniques to detect local translation in neurons are FUNCAT (<u>F</u>l<u>U</u>orescent <u>N</u>on–<u>C</u>anonical <u>A</u>mino acid <u>T</u>agging) and SUnSET (<u>SU</u>rface <u>SE</u>nsing of Translation). The first utilizes modified amino acids, such azidohomoalanine, that get incorporated into the nascent polypeptide chain. The noncanonical amino acids are then tagged with a fluorophore in a cycloaddition reaction (Dieterich et al., 2010). SUnSET is based in the use of the antibiotic puromycin, which mimics an aminoacyl-transferent RNA (tRNA). Puromycin binds to the acceptor site of the ribosome during translation elongation leading to translation termination. The truncated puromycilated polypeptide can be detected by immunofluorescence using an anti-puromycin antibody (Schmidt et al., 2009). The fluorescence signal measured by both approaches is used as a readout of protein synthesis. Nevertheless, the low amount of locally produced proteins entails a limitation in the study of this phenomenon. For instance, our own results indicate that levels of newly-synthesized neuritic proteins can be 20 to 30 times lower than somatic protein levels in unstimulated conditions. Thus, local translation events in neurites can be easily overlooked when analyzing *de novo* synthesis by fluorescence microscopy. To overcome this situation, we have developed a simple method that helps visualize and quantify puromycin-positive sites in neurites by filtering and binarizing imaged cells using FIJI/ImageJ. Moreover, we have used a combination of RNA and protein staining techniques followed by object-based colocalization to detect sites of local RNA translation in neurons.

## Material and Methods

### Animals

All animal protocols followed the European directive 2010/63/EU and were approved by the UPV/EHU ethics committee. Sprague-Dawley rats were bred in local facilities and embryonic brains were obtained from CO_2_ euthanized pregnant rats.

### Neuronal cultures

Hippocampal neurons were prepared from embryonic day 18 rat embryos (E18) as described (Banker and Goslin, 1998). Briefly, hippocampi were dissected from embryonic brains and dissociated in TrypLE Express (Gibco, Thermo Fisher Scientific, Waltham MA, USA) for 10 min at 37°C. Cells were washed twice with Hank’s balanced salt solution (HBSS, Gibco) and resuspended in plating medium containing 10% fetal bovine serum, 2 mM L-glutamine and 50 U.ml^-1^ penicillin-streptomycin in Neurobasal (all from Gibco). Cells were homogenized with a pasteur pipette and centrifuged for 5 minutes at 800 rpm. Cells were resuspended in plating medium. Hippocampal neurons were cultured on poly-D-lysine-coated coverslips in 24-well plates at a density of 20.000 cells/well. Cultures were maintained at 37°C in a 5% CO_2_ humidified incubator. After 1 day *in vitro* (1 DIV) the medium was replaced with growth medium (1X B27, 2 mM glutamine, and 50 U.ml-1 penicillin-streptomycin in Neurobasal). To avoid the growth of glia, half of the medium was replaced with fresh medium containing 20 μM of 5-fluorodeoxyuridine and uridine (Sigma Aldrich, Merck, Darmstadt, Germany) every 3 days. Treatments were performed at 9-10 DIV.

### Oligomeric Aβ preparation and treatments

Soluble oligomeric amyloid-β (Aβ_1-42_) was prepared as previously described (Quintela-Lopez et al., 2019). Synthetic Aβ_1-42_ (Bachem, Bubendorf, Switzerland) was disolved in hexafluoroisopropanol (HFIP, Sigma Aldrich) to 1 mM, aliquoted and dried. For oligomer formation, the peptides were resuspended in dry dimethylsulfoxide (DMSO; 5 mM, Sigma Aldrich) and Hams F-12 (PromoCell Labclinics, Barcelona, Spain) was added to adjust the final concentration to 100 μM. Peptides were incubated overnight at 4°C. Oligomerized Aβ was added to neurons at 9 DIV at a 3 μM concentration and incubated for 24 hours. DMSO was used as vehicle control.

### Puromycylation assay

Puromicyn is a tRNA analog, which is incorporated into the nascent polypeptide chain in a ribosome-catalyzed reaction. This technique allows the *in situ* detection of protein synthesis with an anti-puromicyn antibody. At 10 DIV, DMSO- and Aβ-treated neurons were exposed to 2 μM puromycin (Sigma Aldrich) for 5-30 minutes as indicated. Control conditions with no puromicyn received only fresh growth medium (vehicle). To verify that puromycin labels newly synthesized proteins, 40 μM of the translation inhibitor anisomycin (Sigma Aldrich) was co-incubated with puromycin. Cells were washed with cold PBS with 3 μg.ml^-1^ digitonin (Sigma Aldrich) and fixed in 4% paraformaldehyde (PFA), 4% sucrose in PBS.

### Immunocitochemistry

Neurons were fixed for 20 min at room temperature in 4% PFA, 4% sucrose in PBS. Cells were washed three times with PBS, permeabilized and blocked for 30 min in 3% BSA, 100 mM glycine and 0.25% Triton X-100. Next, samples were incubated overnight at 4°C with primary antibodies against puromycin (1:500, MABE343, Merck Millipore), βIII tubulin (1:500, ab18207 or ab107216, Abcam, Cambridge, UK), Tau (1:1000, ab32057, Abcam) and calreticulin (1:500, ab92516, Abcam). After three PBS washes, cells were incubated for 1 hr at room temperature with fluorophore-conjugated secondary antibodies: anti-mouse Alexa Fluor 594 (1:200, A-11005, Invitrogen, Thermo Fisher Scientific), anti-rabbit Alexa Fluor 488 (1:200, A-21206, Invitrogen), anti-chicken DyLight 350 (1:200, SA5-10069, Invitrogen), anti-rabbit Alexa Fluor 647 (1:200, A-31573, Invitrogen) and anti-rabbit DyLight 405 (1:200, 611-146-002, Rockland Immunochemicals, Pottstown, PA, USA). Samples were washed three times with PBS and mounted with ProLong Gold antifade reagent (P-36930, Invitrogen). Whenever stated, a no-primary-antibody negative control was used.

To label endogenous RNAs neurons were washed once with cold PBS with 3 μg.ml-1 digitonin (Sigma Aldrich), once with 50% methanol in PBS and fixed in cold 100% methanol for 5 minutes. Samples were rehydrated by washing them in 50% methanol in PBS once and in PBS three times. Following the standard immunocytochemistry procedure, cells were incubated for 20 min at room temperature with SYTO RNASelect green fluorescent dye in PBS (1:10.000, S-32703, Invitrogen). Samples were washed with PBS and mounted with ProLong Gold antifade reagent. Some fixed neurons were incubated with 50 μg/ml DNAse or RNAse (Sigma) for 10 minutes at room temperature to assess the selectivity of the SYTO labeling.

### Image acquisition and processing

Images were acquired using a 40x oil objective on an Axio-Observer Z1 microscope equipped with AxioCam MRm Rev. 3 (Zeiss, Oberkochen, Germany) and Hamamatsu CCD (Hamamatsu Photonics, Hamamatsu, Japan) digital cameras. Settings for image acquisition where determined in a random field of a control sample ensuring pixel intensities were within the linear range and avoiding pixel saturation. Settings were kept identical for all sampled cells in any given experiment. Whenever possible, five random cells per coverslip and two coverslips per experimental condition were imaged. Most images were acquired with AxioCam, however if cells were imaged in the far red spectrum, the Hamamatsu camera was used.

For figure preparation, the staining of interest (puromycin, calreticulin, SYTO) was converted from greyscale to RGB or to a colorimetric scale (heatmaps) in non-binarized images. Binarized images used for assisted quantification of translation sites were obtained as will be specified below. In all cases background, contrast and sharpness were adjusted and set the same in control and experimental conditions. Markers used as counterstain for neurite selection were adjusted for an optimal visualization in figures.

### Puromycin and SYTO intensity analysis in non-binarized images

To quantify the puromycin and SYTO fluorescent intensities as measures of protein production and RNA levels respectively, the longest βIII tubulin- and puromycin-positive neurite or Tau- and puromycin-positive neurite from randomly selected cells was straightened with the Segmented Line tool in FIJI/ImageJ:

FIJI/ImageJ > File > Open (do not autoscale) > Segmented Line > Selection > Straighten (line width: 20 pixels for Hamamatsu images; 40 pixels for AxioCam images).

Concentric circles at 10 μm intervals emerging from the center of the cell nucleus or from the edge of the soma were generated with an in-house designed FIJI/ImageJ macro (Concentric_Circles) (Quintela-López et al, 2019). 15 bins were generated covering a length of 150 μm of the straighten neurites. Fluorescence intensity was measured in each bin. Background pixel intensity was measured outside the area covered by the neurite and substracted. To calculate the total fluorescent intensity in the soma or in neurites disregarding the bin position, values retrieved from each bin of interest were summed up.

### Visual inspection of puromycin-positive translation events in non-binarized images (manual analyses)

As described above, the longest positive neurite from randomly selected cells was straightened and divided into 15 10 μm-wide bins with the Concentric_Circles plugin. Discrete puromycin puncta were visually scored in each bin covering a distance of 150 μm from the center of the cell nucleus or from the edge of the soma. To calculate the total translation events in the soma or in neurites disregarding the bin position, values retrieved from each bin of interest were summed up. The bin ranging from 0 to 10 μm (first bin within the soma) was discarded as no discrete puncta could be visualized (N/A in Fig 3D and E).

### Assisted analyses of puromycin- and SYTO-positive events in binarized images

The assisted analysis of translation sites was performed using the following step-by-step protocol:

FIJI/ImageJ > File > Open (do not autoscale). Go to the staining of interest (e.g puromycin) > Process > Filter > Convolve (if a stack is opened, do not process all the images in the stack). The default normalized kernel is sufficient to enhance structures in the periphery of the neurons smaller than 5×5 pixels and it is thus suitable to highlight puromycin-positive translation events distal to the center of the cell nucleus.

Following image convolution:

Image > Adjust > Brightness/Contrast (equal adjustment in all samples within the same experiment) > Image > Type > 8-bit > Process > Binary > Make Binary (Method, MaxEntropy; Background, Default; Black background. Only convert current image).

Once the image is binarized select the longest positive neurite:

Segmented Line > Selection > Straighten (line width: 20 pixels for Hamamatsu images; 40 pixels for AxioCam images) > Process > Smooth > Process > Binary > Make binary (Method, MaxEntropy).

Straighten neurites are finally divided in 15 concentric circles at 10 μm intervals emerging from the center of the cell nucleus or from the edge of the soma with the Concentric_Circles plugin. The number of objects (translation events…) are scored in each interval (bin) with the Analyze Particles function (default settings).

Although this procedure is described for the puromycin staining as an example, the same steps were followed to binarize and quantify SYTO-positive discrete puncta (RNA) in neurites. When binarization of puromycin and SYTO labeling was performed for the same neurite, colocalization between RNA and protein was performed as follows:

Process > Image Calculator > Image 1 (e.g puromycin) AND Image 2 (e.g SYTO) (click create new window).

The resulting image is smoothen and binarized with the MaskEntropy mask. The image is finally divided in 15 concentric circles at 10 μm intervals emerging from the edge of the soma with the Concentric_Circles plugin. The number of objects (considered actively translating RNAs) are scored in each interval (bin) with the Analyze Particles function (default settings).

### Intensity profiles

We used intensity profiles to exemplify how translation events are measured in neurites or to determine the limit of the endoplasmic reticulum (ER) within the somatic region by calreticulin immunostaining. Briefly, cells bodies and/or neurites were selected with the Segmented Line tool (line width: 20 pixels for Hamamatsu images; 40 pixels for AxioCam images) and analyzed with Plot Profile. In neurons stained for calreticulin, the average profile of 5-10 cells per experiment cultured in three independent experiments is shown (Fig 2C).

### Statistical analysis

The sample size is specified in the figure legends. Statistical analyses were performed with Prism 7 (GraphPad Software, San Diego, CA, USA) following a randomized block design where samples from the same experiment were matched to eliminate inter-experimental variability. When comparing the means of two groups taking one variable into account, two-tailed t tests were performed. When comparing the means of two groups taking two variables into account, two-way ANOVA was used. If more than two groups and more than one variable were analyzed, we performed two-way ANOVA followed by Tukey’s multiple comparison test.

For correlation analyses we performed a normality test on the data to determine if they followed a Gaussian distribution, which most of them didn’t. Thus, we chose to perform Spearman nonparametric correlation test to retrieve the correlation coefficients. In the correlation graphs, linear regression of the data was performed to evaluate the differences between slopes (ANCOVA).

To evaluate the calreticulin staining, the background from the no-primary antibody negative control was subtracted and a one-sample t test was performed.

In all tests p < 0.05 was considered statistically significant.

## Results

### Detection of newly-synthesized neuritic proteins by puromycilation

Based on previously published data (Baleriola et al., 2014), rat hippocampal neurons grown for 9 days *in vitro* (DIV) were treated with vehicle (DMSO) or 3 μM Aβ_1-42_ oligomers for 24 hours by bath application. As a first step to quantify RNA translation sites in neurites we first detected *de novo* production of neuritic proteins by puromycilation/SUnSET (Schmidt et al., 2009) (Fig 1). The antibiotic puromycin is an aminoacyl-tRNA analog that incorporates into the polypeptide chains during translation elongation, leading to translation termination (Yarmolinsky and Haba, 1959). The development of specific antibodies has allowed the immunodetection of puromycilated polypetides as a measure of protein synthesis. Control and Aβ-treated cells were fed with 2 μM puromycin for 30 minutes prior to fixation. Following fixation with a PFA / sucrose mix, cells were stained for puromycin and counterstained with an anti-βIII tubulin antibody to visualize the neuronal cytoskeleton (Fig 1A). As a negative control, immunostaining was performed on neurons that had not been treated with puromycin (-puro, Fig 1A). Fluorescence intensity for the raw puromycin signal, represented either in red (RGB, Fig 1A) or in a colorimetric scale (heatmap, Fig 1A), was measured along the longest positive neurite in randomly selected cells (1-6, Fig 1A). Fluorescence levels in puromycin-labeled neurites (3 and 4, Fig 1A-C) were well above the levels measured in negative controls (1 and 2, Fig 1A-C). Additionally, puromycin hotspots were readily visible in distal sites of the neurites, especially in Aβ-treated cells (4, intensity profile and heatmap in Fig 1B). Finally, regardless of the effect of Aβ_1-42_, puromycin intensity was significantly reduced in neurites when cells were co-incubated with the translation inhibitor anisomycin (+anis+puro, Fig 1A; 5 and 6, Fig 1A-C). Altogether these results indicate that in our system puromycin labeling can be used to detect *de novo* synthesis of neuritic proteins as previously reported in similar experimental setups (Walker et al.,2018; Rangaraju et al., 2019).

**Figure 1.**
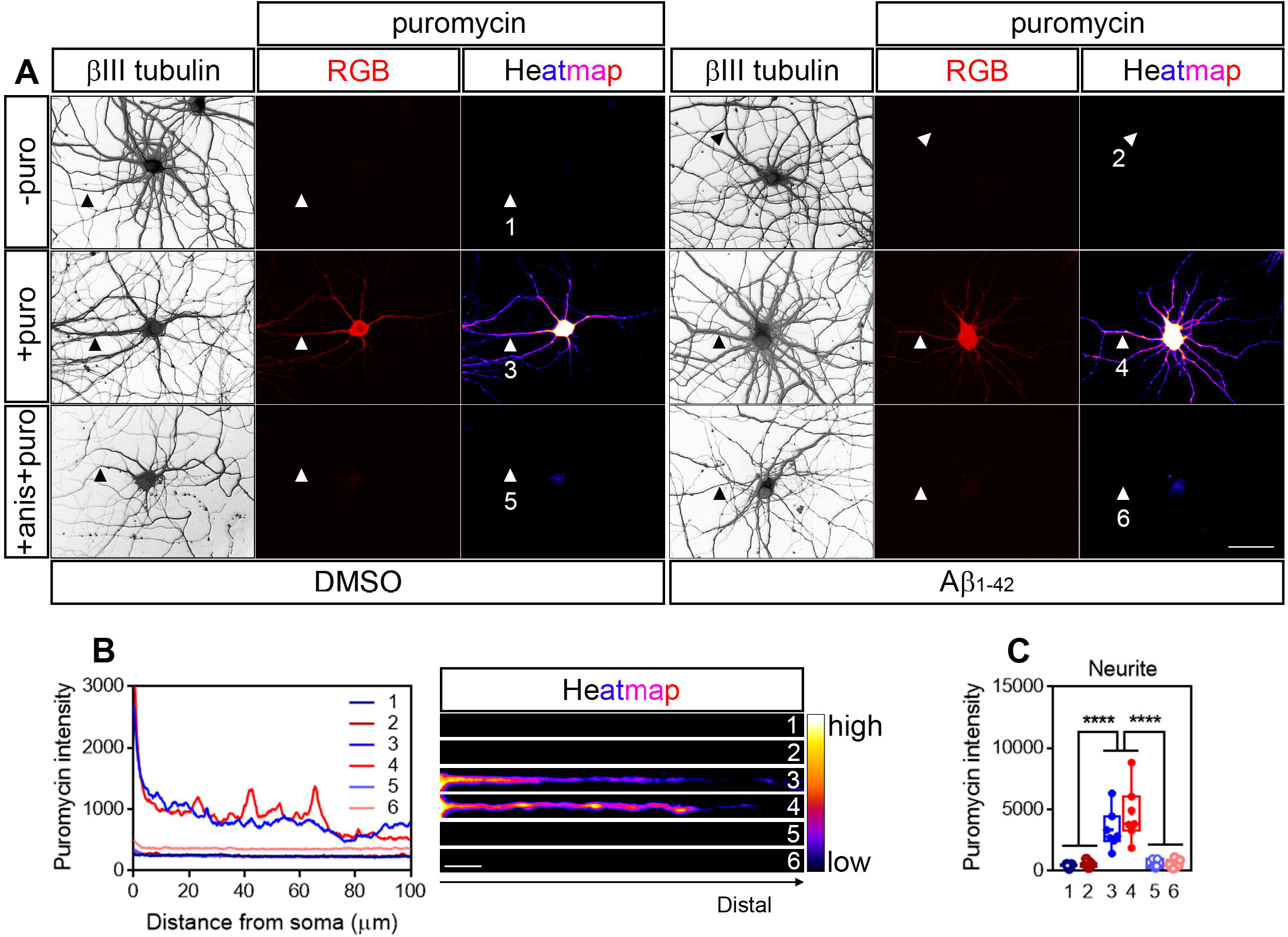
Detection of newly synthesized proteins by puromycilation. **(A)** Rat hippocampal neurons were grown for 9 DIV and were treated with DMSO (left panels) or Aβ_1-42_ oligomers (right panels) for 24 h. Before fixing, cells were incubated with vehicle (-puro; neurites 1 and 2), with puromycin (+puro; neurites 3 and 4) or with puromycin and anisomycin (+anis+puro; neurites 5 and 6) for 30 mins. Cells were immunostained with an anti-βIII tubulin antibody to visualize the neuronal cytoskeleton (grey) and with an anti-puromycin antibody to analyze newly-synthesized proteins (RGB and heatmaps). Scale bar, 50 μm. **(B)** Intensity profiles were measured in the longest positive neurite from randomly selected cells as exemplified. 1 and 2; no puromycin incubation in DMSO- and Aβ-treated neurons respectively. 3 and 4; 30 min puromycin incubation in DMSO- and Aβ-treated cells respectively. 5 and 6; co-incubation with anisomycin and puromycin for 30 mins in DMSO- and Aβ-treated cells respectively. Scale bar, 10 μm in heatmaps. **(C)** Box and whisker graph representing the total fluorescent intensity of the puromycin staining along 130 μm of βIII tubulin-positive neurites. 1 and 2; no puromycin incubation in DMSO- and Aβ-treated neurons respectively. 3 and 4; 30 min puromycin incubation in DMSO- and Aβ-treated cells respectively. 5 and 6; co-incubation with anisomycin and puromycin for 30 mins in DMSO- and Aβ-treated cells respectively. Data represent the average value of 5-10 sampled cells per condition shown as individual data points, and the mean and median of 7 independent experiments (n=7). **** p < 0.0001; two-way ANOVA followed by Tukey’s multiple comparison test.

We then asked whether Aβ oligomers induced changes in the distribution pattern of newly synthesized proteins along neurites. The longest puromcycin-positive neurite of randomly selected cells was straighten and divided into 10 μm bins. Puromycin intensity was measured in 15 bins covering a length of 150 μm from the center of the cell nucleus using the Concentric_Circles plugin in FIJI/ImageJ (Fig 2A). A significantly distinct distribution in the levels of newly produced proteins was observed in Aβ-treated neurites compared to controls (positions beyond 20 μm, Fig 2B). Then we asked whether the puromycin signal likely arose from the endoplasmic reticulum (ER). Soma-centric views consider that most protein synthesis in eukaryotic cells occurs in the ER (specifically in the rough ER). In neurons, however, the positioning of the rough ER (RER) with respect to distal sites of neurites does not explain how in some experimental setups, that allow to study the local response of dendrites and axons (reviewed in (Holt et al., 2019) newly synthesized proteins are detected peripherally, unless they are produced locally beyond the “canonical” ER. The RER is enriched in proteins involved in the folding of nascent polypeptides, being the Calnexin/Calreticulin system one the best known protein complex (Rutkevich and Williams, 2011). After culturing hippocampal neurons for 10 days, cells were processed for Calreticulin (Calr) immunostaing. As a negative control, some neurons were subjected to the immunocytochemistry procedure but were not incubated with anti-Calr antibody (no-primary antibody control). To determine the extension of the Calr staining in our neurons we substracted the fluorescent signal from cells incubated without a primary antibody to those incubated with anti-Calr. We determined that neuritic positions 50 μm beyond the nucleus were devoid of Calreticulin, and thus of “canonical” ER (Fig 2C). Interestingly, Aβ significantly increased the levels of newly synthesized proteins in the 50-150 μm interval (distal, Fig 2A, C and D). More importantly the effect of Aβ was restricted to neurites and did not affect the neuronal soma (Fig. 2D).

**Figure 2.**
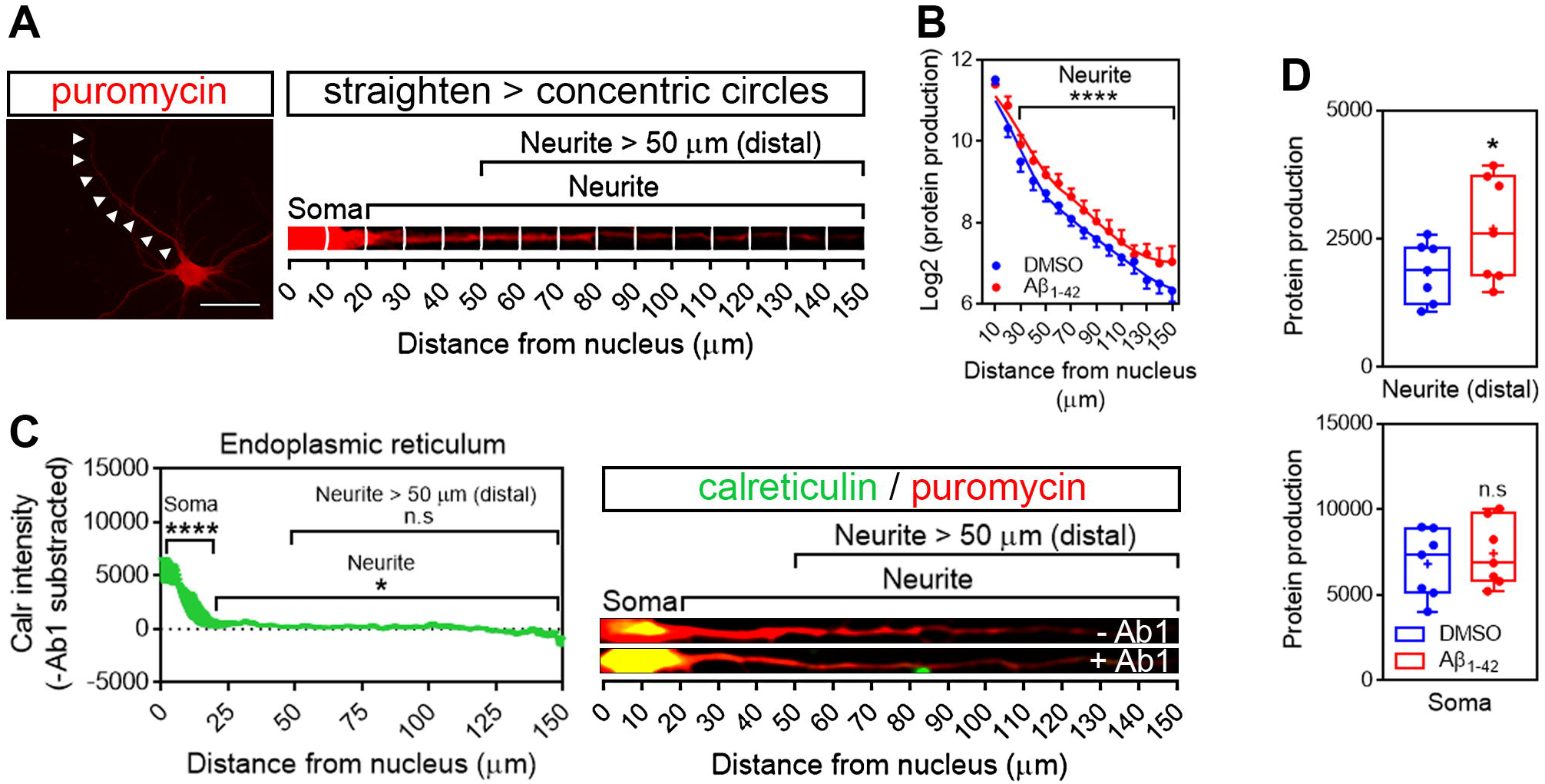
Aβ_1-42_ oligomers increase *de novo* synthesis of neuritic proteins. **(A)** Rat hippocampal neurons were grown for 9 DIV and treated with DMSO or with Aβ_1-42_ oligomers for 24 h. Cells were processed for puromycin staining to measure protein synthesis (red) and counterstained with an anti-βIII tubulin antibody to visualize the neuronal cytoskeleton (not shown). The longest positive neurite (arrowheads in the red puromycin micrograph) was selected with a segmented line, straighten and divided into 10 μm bins with the Concentric_Circles plugin (right). Scale bar, 50 μm. **(B)** Puromycin intensity was measured in DMSO- and Aβ-treated neurons in 15 bins covering a distance of 150 μm from the cell nucleus (as also shown in the straighten micrograph exemplified in **(A)**). The graph shows the average intensity of puromycin per condition represented as Log2(protein production) vs distance +/- SEM measured in 7 independent experiments (n=7). **** p < 0.0001; two-way ANOVA. **(C)** The “canonical” endoplasmic reticulum was defined by calreticulin staining (Carl in green). To determine the background signal, some cells were stained only with the secondary antibody (no-primary antibody control). Background signal from the no-primary antibody control (-Ab1) was subtracted from the actual anti-calreticulin antibody signal (+Ab1). The intensity profile graph represents the average calreticulin signal from DMSO- and Aβ-treated cells, after background subtraction, cultured in 3 independent experiments (n=3). **** p < 0.0001; * p < 0.05; n.s, no significant; one-sample t test. **(D)** Box and whisker graphs representing the total fluorescent intensity of the puromycin staining in βIII tubulin-positive neurites within the range of 50-150 μm from the nucleus (Neurite (distal)) as also exemplified in **(A)**) and in the soma (soma; 0-20 μm as also exemplified in **(A)**). Data represent the average value of 5-10 sampled cells per condition shown as individual data points, and the mean and median of 7 independent experiments (n=7). * p < 0.05; two-tailed t test.

### Image processing unravels a previously unreported effect of Aβ_1-42_ oligomers on translation hotspots in neurites

Measuring puromycin intensity can give an idea of the amount of protein being produced distal from the ER within neurites and/or diffused from the actual translation site, but it does not report on the number and position of the translation sites themselves. In the case of Aβ treated cells, increased puromycin intensity might be a result of the emergence of new translation sites, a consequence of an increased rate of protein production in preexisting sites or both. To determine whether Aβ oligomers modify the amount of translation sites or events in neurites we quantified the number of puromycin discrete puncta. Discrete puncta in distal neuritic sites likely reflect foci of localized translation (Graber et al., 2013; Rangaraju et al., 2019). Such foci can be easily overlooked since their intensity can be ~20 to 30 times less than somatic puromycin fluorescent levels (as implicitly shown in Fig 2). We have developed a strategy to enhance puromycin hotspots in neurites based solely on image processing and the assisted quantification of the resulting objects.

The number of discrete puromycin foci were quantified along the longest puromycin-positive neurite of randomly sampled cells (Fig 3A). Image acquisition was identical in control and Aβ-treated neurons. We selected neurites from raw and binarized images in order to compare quantifications performed by visual inspection of the puromycin staining (manual) and by analyzing particles (assisted) respectively. On the one hand DMSO- and Aβ-treated neurites (1 and 2, Fig 3B) were selected from raw images (represented as RGBs or heatmaps in Fig 3B) with a segmented line 20 or 40 pixels wide and straighten. Next, the same images were filtered with the convolver in FIJI/ImageJ applying the default normalized kernel. The default kernel was sufficient to enhance structures in the periphery of the neurons and thus was suitable to highlight puromycin-positive translation events distal to the center of the cell nucleus.

**Figure 3.**
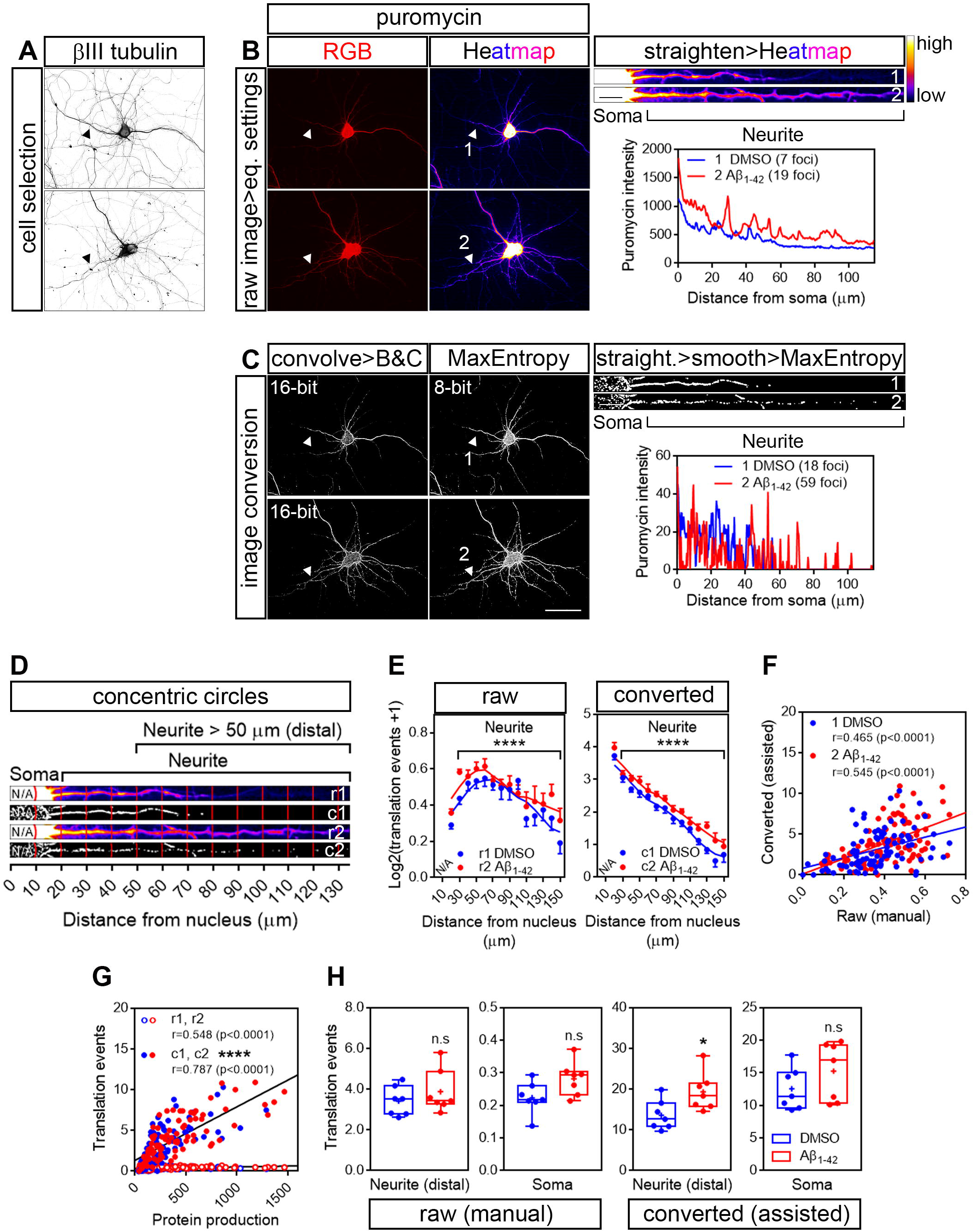
Image processing reveals an effect of Aβ_1-42_ oligomers on neuritic translation sites. **(A)** Rat hippocampal neurons were grown for 9 DIV and were treated with DMSO or Aβ_1-42_ oligomers for 24 h. Cells were incubated with puromycin for 30 mins and processed for βIII tubulin (grey) and **(B)** puromycin immunostaining (RGB and heatmaps). The longest puromycin-positive neurite was selected with a segmented line and straighten. Puromycin-positive discrete puncta were analyzed by visual inspection as exemplified in the intensity profiles obtained from straighten neurites (heatmaps). Scale bar, 50 μm in **(A)**, and in RGB and heatmaps in **(B)**; 10 μm in straighten neurites (heatmap) in **(B)**. **(C)** Puromycin staining from randomly selected cells was filtered with the convolver applying the default normalized kernel and brightness and contrast were adjusted (convolve>B&C). 16-bit images were coverted to 8-bit and binarized with the MaxEntropy mask (MaxEntropy). Neurites were selected with a segmented line, straighten, smoothen and binarized with the MaxEntropy mask (straight>smooth>MaxEntropy). Puromycin-positive discrete puncta were analyzed with the particle analyzer as exemplified in the intensity profiles from straighten neurites. Scale bar, 50 μm in converted images; 10 μm in converted straighten neurites. **(D)** Discrete puromycin puncta were measured in DMSO- and Aβ-treated neurons in 15 bins covering a distance of 150 μm from the cell nucleus using the Concentric_Circles plugin. Measurements were performed by visual inspection in raw images (r1, DMSO-; r2, Aβ-treated neurites) and with the particle analyzer in binarized images (c1, DMSO-; c2, Aβ-treated neurites). N/A, not applicable. **(E)** Graphs show the average puromycin puncta per condition represented as Log2(translation events+1) vs distance +/- SEM measured in 7 independent experiments (n=7). **** p < 0.0001; two-way ANOVA. N/A, not applicable. **(F)** Spearman correlation between quantifications in raw and in binarized images. Graphs represent each scored value per distance using both methods in DMSO- (blue) and Aβ-treated neurons (red) cultured in 7 independent experiments (n=7). p < 0.05 indicate a significant correlation. **(G)** Spearman correlation between non-assisted (r1, DMSO-; r2, Aβ-treated cells) or assisted quantification (c1, DMSO-; c2, Aβ-treated neurons) of translation events and puromycin intensity (represented as protein production). Graphs represent each scored value per distance from 7 independent experiments (n=7). p < 0.05 indicate a significant correlation. **** p < 0.0001; significant differences between slopes. **(H)** Box and whisker graphs representing the total translation events in βIII tubulin-positive neurites within the range of 50 to 150 μm (Neurite (distal)) and in the soma (soma; 0-20) following visual inspection of raw images (raw (manual)) or assisted quantification in binarized images (converted (assisted)). Data represent the average value of 5-10 sampled cells per condition shown as individual data points, and the mean and median of 7 independent experiments (n=7). * p < 0.05; n.s, no significant; two-tailed t test.

Brightness and contrast were then manually adjusted in order to eliminate pixels outside the stained cells (background). 16-bit images were coverted to 8-bit and binarization was performed using the MaxEntropy mask in FIJI/ImageJ (Figure 3C). As in the case of the raw images, DMSO- and Aβ-treated neurites (1 and 2, Fig 3C) were selected with a 20- or 40-pixel wide segmented line and straighten. Finally, straighten neurites were smoothen and binarized again using the MaxEntropy mask. As shown by the number of peaks in the intensity profiles (3B and C), image conversion increased the number of detected events (foci in 3B and C) and slightly enhanced the effect of Aβ oligomers, which increased from 2.7-to 3.3-fold. We then analyzed the distribution pattern of translation foci along neurites. Neurites were divided into 10 μm bins and positive puromycin puncta within each bin were visually scored prior to image conversion (r1 and r2, Fig 3D) or were counted with the particle analyzer in binarized images (c1 and c2, Fig 3D). In all cases 15 bins were quantified per cell, covering a distance of 150 μm from the cell nucleus. Both quantification methods revealed a significantly distinct distribution of translation events in neurites in Aβ-treated cells compared to controls (Fig 3E). When comparing the scores performed at each distance by manual inspection in raw images and with the assisted method in binarized images we observed a significant positive correlation between both procedures that ranged from moderate to high in DMSO- and Aβ-treated cells respectively (Fig 3F). To determine which method was closer to the unbiased measurement of protein production represented by puromycin intensity (Fig 2),we then compared data obtained from binarized images and from raw images with the intensity values. In both cases we found a significant high positive correlation (Fig 3G). However, when fitting the translation events at each distance to a regression line, a significant increase in the slope was observed when data were obtained from binarized images, suggesting increased similarities between the number of translation sites and the amount of protein produced when using the assisted quantification method. Finally, we focused on distal sites of the neurites (> 50 μm from the nucleus) disregarding the bin position and were unable to detect any significant change between DMSO- or Aβ-treated cells when translation events were quantified in raw images by visual inspection (manual, Fig 3H). Conversely, we did observe a significant effect of Aβ oligomers when quantification was performed with the particle analyzer in binarized images (assisted, Fig 3H). In no case did we detect any changes induced by Aβ in the soma. Altogether, results so far indicate that binarizing images from puromycin-positive cells allows the assisted quantification of neuritic translation sites yielding results that resemble those obtained from an unbiased measurement of raw puromycin intensity. Additionally, assisted quantification in binarized images enhances the effect of Aβ oligomers on discrete puromycin puncta in distal neurites.

The first evidence of Aβ oligomers regulating local translation in neurons was reported in axons (Baleriola et al., 2014). Thus, we next tried our assisted quantification method in neurites positive for the axonal protein Tau. Additionally, after treatment with DMSO or Aβ oligomers for 24 hours, we fed the cells with puromycin for 5, 10 or 30 minutes. Puromycin pulses shorter than 30 minutes have been successfully used to detect localized translation in neurites in other experimental setups (Graber et al., 2013; Walker et al.,2018; Rangaraju et al., 2019). We first analyzed the distribution pattern of protein production along Tau-positive neurites. The longest positive neurite was selected from randomly sampled cells imaged with identical settings. Neurites from raw images (exemplified as heatmaps in Fig 4A) were straighten and divided into 10 μm bins. Puromycin intensity was measured in 15 bins covering a distance of 150 μm from the edge of the soma (Fig 4B). No changes in newly synthesized proteins were observed between control and Aβ-treated cells when neurons were exposed to puromycin for 5 or 10 minutes. However, a significantly distinct pattern in protein production induced by Aβ oligomers was detected in Tau-positive neurites following a 30-minute treatment with puromycin (Fig 4B). Focusing on distal sites of the neurite (beyond 30 μm from the soma) we observed a significant accumulation of newly synthesized proteins after 30 minutes of puromycin treatment compared to shorter exposures in both DMSO- and Aβ-treated cells. A significant increase in protein production in Aβ-treated neurites compared to controls was also detected with the longest puromycin exposure (Fig 4C). These results, similar to the ones obtained in βIII tubulin-positive neurites, confirm that Aβ oligomers induce *de novo* synthesis of axonal proteins as previously reported (Baleriola et al., 2014;Walker et al., 2018). We then quantified the number of translation events. Images were convolved with the default normalized kernel in FIJI/ImageJ and processed like βIII tubulin neurites as described before. Translation events were scored with the particle analyzer in 15 bins covering a distance of 150 μm from the edge of the cell body. Again, a distinct pattern of translation was observed between DMSO- and Aβ-treated neurites only when cells were fed with puromycin for 30 minutes (Fig 4E). Similarly, despite detecting a significant accumulation of translation events in both control and Aβ-treated cells after 30 minutes of puromycin exposure compared to shorter pulses, these were significantly higher when Aβ oligomers were added to the cultures (Fig 4F).

**Figure 4.**
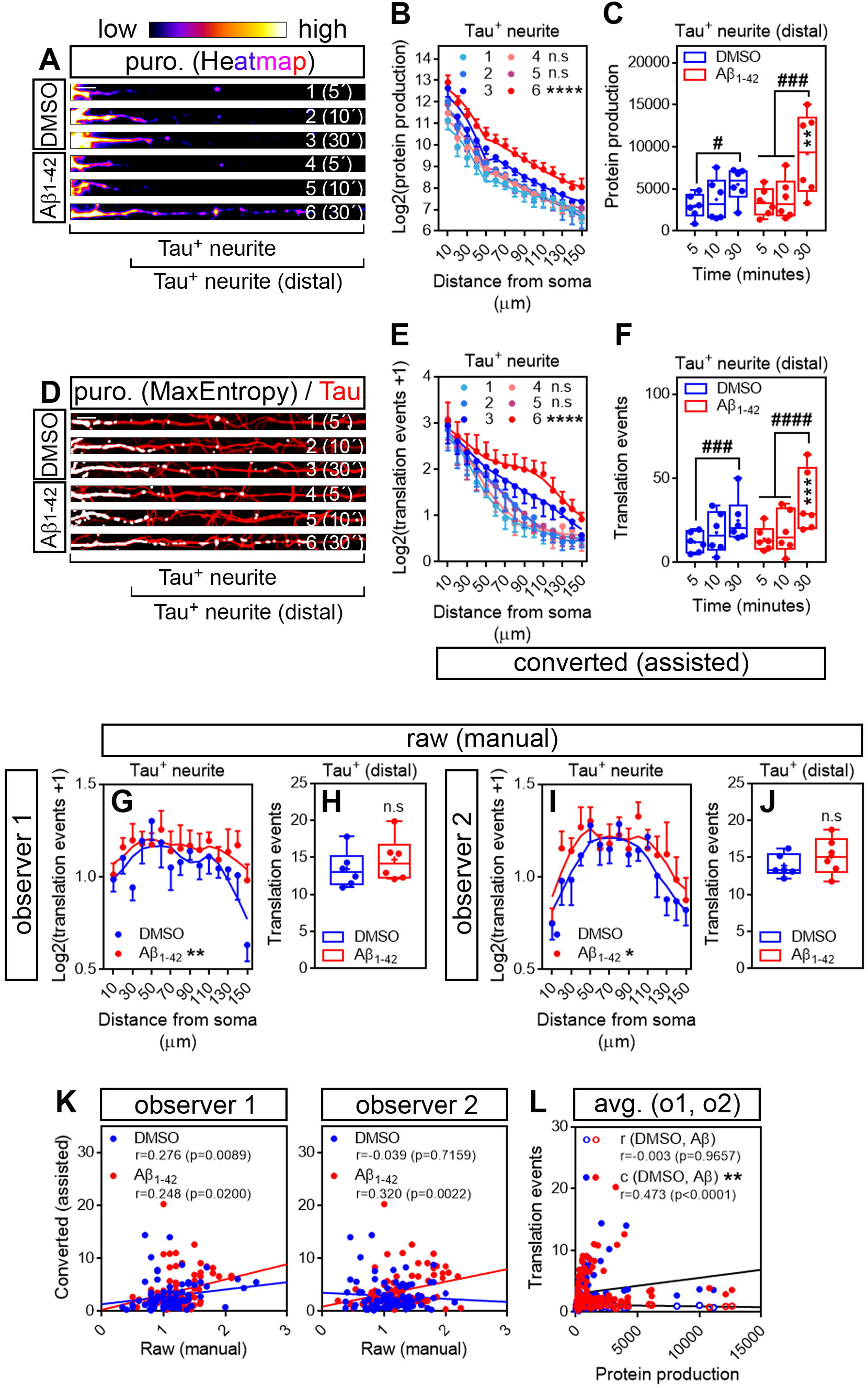
Aβ_1-42_ oligomers increase translation sites in Tau-positive neurites. **(A)** Rat hippocampal neurons were grown for 9 DIV and treated with DMSO or with Aβ_1-42_ oligomers for 24 h. Cells were fed with puromycin for 5, 10 or 30 mins, fixed and immunostained with an anti-puromycin antibody to measure protein synthesis (heatmaps) and counterstained with an anti-Tau antibody (not shown). The longest Tau-positive neurite was selected with a segmented line and straighten. 1, 2, and 3, DMSO-treated cells exposed to puromycin for 5, 10 and 30 mins respectively; 4, 5 and 6, Aβ-treated cells exposed to puromycin for 5, 10 and 30 mins respectively. Scale bar, 10 μm. **(B)** Puromycin intensity was measured in DMSO- and Aβ-treated neurons in 15 bins covering a distance of 150 μm from the edge of the soma. The graph shows the average intensity of puromycin per condition represented as Log2(protein production) vs distance +/- SEM measured in 6 independent experiments (n=6). 1, 2, and 3, DMSO-treated cells exposed to puromycin for 5, 10 and 30 mins respectively; 4, 5 and 6, Aβ-treated cells exposed to puromycin for 5, 10 and 30 mins respectively. **** p < 0.0001 DMSO vs Aβ, 30 mins puromycin; two-way ANOVA followed by Tukey’s multiple comparison test. **(C)** Box and whisker graphs representing the total fluorescent intensity of the puromycin staining in Tau-positive neurites within the range of 30 to 150 μm (Tau^+^ (distal) as exemplified in **(A)**). Data represent the average value of 10-20 sampled cells per condition shown as individual data points, and the mean and median of 6 independent experiments (n=6). # p < 0.05 5 vs 30 mins puromycin in DMSO-treated cells; ### p < 0.001 5 vs 30 mins and 10 vs 30 mins in Aβ-treated neurons; ** p < 0.01 DMSO vs Aβ, 30 mins puromycin; two-way ANOVA followed by Tukey’s multiple comparison test. **(D)** Puromycin staining from randomly selected cells was filtered with the convolver, brightness and contrast were adjusted. Images were converted to 8-bit and binarized with the MaxEntropy mask. The longest Tau-positive neurite was selected with a segmented line and straighten, smoothen and binarized with the MaxEntropy mask (MaxEntropy). Counterstain with the anti-Tau antibody is shown (red). 1, 2, and 3, DMSO-treated cells exposed to puromycin for 5, 10 and 30 mins respectively; 4, 5 and 6, Aβ-treated cells exposed to puromycin for 5, 10 and 30 mins respectively. Scale bar, 10 μm. **(E)** Puromycin-positive discrete puncta were scored with the particle analyzer in 15 bins covering a distance of 150 μm from the edge of the soma. The graph shows the average translation events per condition represented as Log2(translation events+1) vs distance +/- SEM measured in 6 independent experiments (n=6). 1, 2, and 3, DMSO-treated cells exposed to puromycin for 5, 10 and 30 mins respectively; 4, 5 and 6, Aβ-treated cells exposed to puromycin for 5, 10 and 30 mins respectively. **** p < 0.0001 DMSO vs Aβ, 30 mins puromycin; two-way ANOVA followed by Tukey’s multiple comparison test. **(F)** Box and whisker graphs representing the total number of translation events scored in Tau-positive neurites within the range of 30 to 150 μm (Tau^+^ (distal) as exemplified in **(D)**). Data represent the average value of 10-20 sampled cells per condition shown as individual data points, and the mean and median of 6 independent experiments (n=6). ### p < 0.001 5 vs 30 mins puromycin in DMSO-treated cells; #### p < 0.0001 5 vs 30 mins and 10 vs 30 mins in Aβ-treated neurons; *** p < 0.001 DMSO vs Aβ, 30 mins puromycin; two-way ANOVA followed by Tukey’s multiple comparison test. **(G)** Discrete puromycin puncta scored by observer 1 in DMSO- and Aβ-treated neurons in 15 bins covering a distance of 150 μm from the edge of the soma and in **(H)** distal sites of Tau-positive neurites disregarding the bin position. **(I)** Discrete puromycin puncta scored by observer 2 in DMSO- and Aβ-treated neurons in 15 bins covering a distance of 150 μm from the edge of the soma and in **(J)** distal sites of Tau-positive neurites disregarding the bin position. All measurements were performed by visual inspection in raw images. Graphs in **(G)** and **(I)** show the average number of translation events per condition represented as Log2(translation events+1) vs distance +/- SEM measured following a 30-minute puromycin pulse in 6 independent experiments (n=6). * p < 0.05; ** p < 0.01; two-way ANOVA followed by Tukey’s multiple comparison test. Box and whisker graphs in **(H)** and **(J)** show the total number of translation events scored in Tau-positive neurites within the range of 30 to 150 μm (Tau+ (distal)). Data represent the average value of 10-20 sampled cells per condition shown as individual data points, and the mean and median of 6 independent experiments (n=6). n.s, no significant; two-tailed t tests. **(K)** Spearman correlation between quantifications in raw (manual) and in binarized (assisted) images. Graphs show values scored in raw (manual) images by observer 1 and observer 2 in DMSO- (blue) and Aβ-treated neurons (red) cultured in 6 independent experiments (n=6). p < 0.05 indicate a significant correlation. **(L)** Spearman correlation between non-assisted (r (DMSO, Aβ)) or assisted quantification (c (DMSO, Aβ)) of translation events and puromycin intensity (represented as protein production). Graphs represent the non-assisted counts per distance as the average score obtained by observers 1 and 2. Data correspond to 6 independent experiments (n=6). p < 0.05 indicate a significant correlation. ** p < 0.01; significant differences between slopes.

To determine if our assisted scoring method correlated better than manual quantification with the unbiased measurements of fluorescence intensity also in Tau-positive neurites, two independent observers quantified the number of puromycin-positive puncta along neurites by visual inspection of raw images (Fig 4G-J). Both observers reported a significantly distinct distribution of translation sites in DMSO- and Aβ-treated samples when scores were performed in 10 μm bins (Fig 4G and I). However, when focusing on distal sites of the neurites (> 30 μm from the soma) disregarding the bin position, none of them detected changes between controls and Aβ treatments (Fig 4H and J), in line with previous results (Fig 3H). Data retrieved from observer 1 revealed a low yet significant correlation between scores obtained in binarized images and those obtained in raw images in both control and Aβ-treated neurons, whereas the correlation between both scoring methods was only significant upon Aβ treatment based on results from observer 2 (Fig 4K). We then compared data obtained from binarized images and the averaged data retrieved from observers 1 and 2 with the intensity values. We found no significant correlation between the amount of protein produced at each neuritic position and the number on translation sites scored by visual inspection (r, Fig 4L). Conversely, a significant moderate positive correlation was observed between parameters when translation sites were counted in binarized images with the particle analyzer (c, Fig 4L). Furthermore, when fitting the translation events at each distance to a regression line, a significant increase in the slope was observed when data were obtained from binarized images, suggesting increased similarities between the number of translation sites and the amount of protein produced when using the assisted quantification method (Fig 4L). These results not only confirm that scoring puromycin-positive sites in neurites in binarized images by assisted means show a better fit with the unbiased measurement of raw puromycin intensity, but also reveal an effect of Aβ oligomers on discrete translation events in neurites that was previously unreported.

### Object-based colocalization analyses confirm that peripheral translation events arise from localized RNAs

As mentioned previously, discrete puromycin-positive puncta in distal neurites likely reflect sites of local translation. Nevertheless, we sought to determine if in our system what we had reported as neuritic translation sites did in fact colocalize with neuritic RNAs. Samples processed for puromycin detection in Tau-positive neurites were incubated with SYTO RNASelect, a fluorescent dye that selectively binds RNA (Savas et al., 2010). To verify that SYTO could be successfully used in our system to label neuritic RNA we compared the fluorescent intensity of the dye within Tau positive neurites to background fluorescent levels in cells that had not been incubated with SYTO. Additionally, some fixed cells were digested with DNAse or RNAse prior to labeling. Neurites from SYTO-positive cells showed significantly higher levels of fluorescence than those not incubated with the dye (graphs and neurites 1 and 2 in Fig 5A). More importantly, levels of SYTO were similar in positive neurites incubated in the presence or absence of DNAse (graphs and neurites 2 and 3 in Fig 5A), whereas incubation with RNAse moderately yet significantly reduced the fluorescence intensity (graphs and neurite 4 in Fig 5A). These results indicate that indeed neuritic RNAs can be labeled with SYTO RNASelect dye.

**Figure 5.**
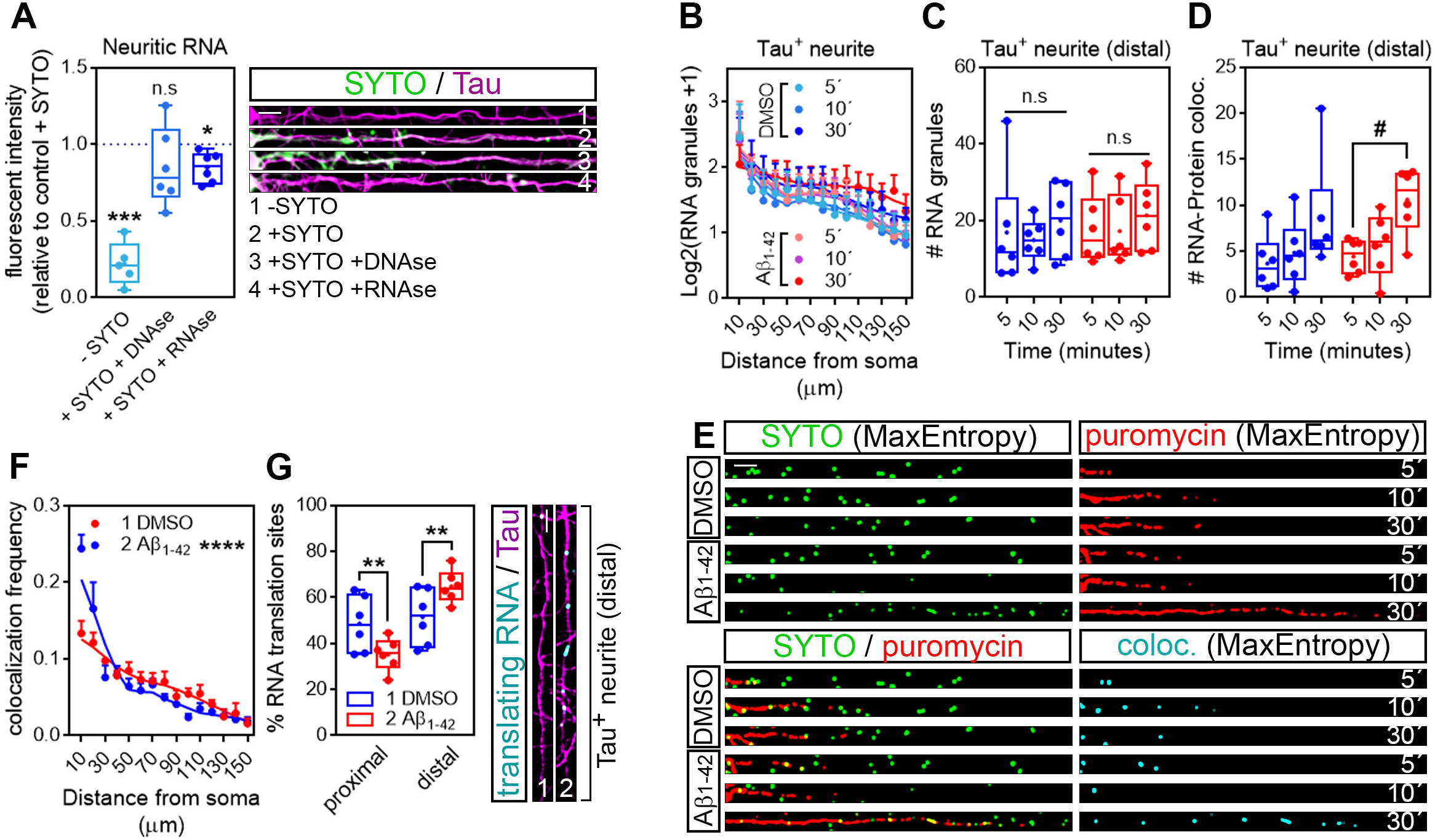
Translation events in Tau-positive neurites are a result of local protein synthesis. **(A)** Cells grown for 9 DIV and treated with DMSO for 24 h. Cells immunostained with an anti-Tau antibody (magenta) were incubated with SYTO RNASelect green fluorescent dye to label endogenous RNA (green). Total green fluorescence intensity was measured in neurites covering a distance of 150 μm from the edge of the soma (2, + SYTO). As negative control, green fluorescence was measured in cells that had not been incubated with SYTO (1, -SYTO). To determine if SYTO selectively labeled RNA, some fixed cells were digested with DNAse (3, +SYTO +DNAse) or with RNAse (4, +SYTO +RNAse). Box and whisker graphs represent the average relative fluorescence intensity of 10 neurites per condition, shown as individual data points, and the mean and median of 5 (n=5, -SYTO negative samples compared to their corresponding +SYTO controls) or 6 (n=6, +SYTO + DNAse and +SYTO +RNAse compared to their corresponding + SYTO controls) independent experiments. *** p < 0.001; * p < 0.05; n.s, not significant; two-tailed t tests. Scale bar, 10 μm. **(B)** SYTO-positive staining (as represented in green in **(E)**) from randomly selected cells was filtered with the convolver, brightness and contrast were adjusted. Images were converted to 8-bit and binarized with the MaxEntropy mask. The longest Tau-positive neurite was selected with a segmented line and straighten, smoothen and binarized with the MaxEntropy mask (MaxEntropy). SYTO-positive discrete puncta were scored with the particle analyzer in 15 bins covering a distance of 150 μm from the edge of the soma. The graph shows the average number on puncta represented as Log2(RNA granules+1) vs distance +/- SEM measured in 6 independent experiments (n=6). No statistical differences were detected between DMSO- and Aβ-treated cells incubated with puromycin for 5, 10 or 30 mins. **(C)** Box and whisker graphs show the total number of RNA granules (SYTO-positive puncta) in distal sites of Tau-positive neurites from DMSO- and Aβ-treated cells incubated with puromycin for 5, 10, or 30 mins. Data represent the average value of 10 sampled cells per condition plotted as individual data points, and the mean and median of 6 independent experiments (n=6). n.s, no significant; two-way ANOVA followed by Tukey’s multiple comparison test. **(D)** Parallel to processing SYTO-labeled images, puromycin staining was filtered with the convolver, brightness and contrast were adjusted. Images were converted to 8-bit and binarized with the MaxEntropy mask. Same Tau-positive neurites selected for SYTO quantification (green channel) were selected in the red channel (puromycin staining in **(E)**), strainghten, smoothen and binarized with the MaxEntropy mask. Co-localized objects were obtained with the AND function in the image calculator (cyan in **(E)**) and scored in distal sites of Tau-positive neurites with the particle analyzer. Box and whisker graphs show the total RNA-protein colocalized puncta in DMSO- and Aβ-treated cells incubated with puromycin for 5, 10, or 30 mins. Data represent the average value of 10 sampled neurites per condition plotted as individual data points, and the mean and median of 6 independent experiments (n=6). # p< 0.05 5 min vs 30 min puromycin in Aβ-treated cells; two-way ANOVA followed by Tukey’s multiple comparison test. **(E)** Micrographs from straighten, binarized neurites stained with SYTO RNASelect green fluorescent dye to label RNA (green), with an anti-puromycin antibody to label protein (red) and the resulting images when merging both channels (green, red and yellow) and when combining both with the AND function in the image calculator (cyan). Cells treated with puromycin for 5, 10 or 30 mins are shown. Scale bar, 10 μm. **(F)** The graph represents the frequency distribution of SYTO- and puromycin-positive objects (colocalization frequency) in DMSO- and Aβ-treated neurites following 30-min puromycin exposure. Colocalized objects were measured with the particle analyzer in 15 bins covering a distance of 150 μm from the edge of the cell body. **** p < 0.0001 (interaction); two-way ANOVA. **(G)** Box and whisker graph representing the proportion of colocalized objects (% RNA translation sites) in proximal (0-30 μm) and distal (last 120 μm) sites of Tau-positive neurites. Data represent the average value of 10 sampled cells per condition shown as individual data points, and the mean and median of 6 independent experiments (n=6). ** p < 0.01; two-tailed t test. Micrographs show colocalized objects (translating RNAs, cyan) detected within the last 120 μm (distal) of Tau positive neurites (magenta). Scale bar, 10 μm.

We then analyzed the distribution of RNA granules, measured as SYTO-stained foci, along Tau-positive neurites. For this purpose, raw images stained for SYTO were processed following the exact same protocol as for puromycin labeling: images were convolved with the default normalized kernel in FIJI/ImageJ, brightness and contrast were manually adjusted, 16-bit images were converted to 8-bit and binarized using the MaxEntropy mask. Neurites were then selected with a segmented line, straighten, smoothen and binarized again with the MaxEntropy function (green, Fig 5E). The number or RNA granules was scored in 15 bins covering a distance of 150 μm from the edge of the soma and no significant differences were observed between experimental conditions, regardless of whether neurons were fed with puromycin for 5, 10 or 30 minutes (Fig 5B). Similarly, no significant changes were detected in distal sites (> 30 μm from the soma) between DMSO- and Aβ-treated neurites (Fig 5C). Thus, Aβ treatment does not affect RNA recruitment to neurites. In these experiments, green and red channels corresponding to RNA (SYTO, Fig 5E) and protein (puromycin, Fig 5E) were binarized in parallel and colocalization between objects in both channels was calculated using the AND function in the FIJI/ImageJ image calculator. The resulting puncta (cyan, Fig 5E) were scored in 10 μm bins covering a distance of 150 μm from the edge of the cell body. No significant differences between DMSO- and Aβ-treated cells were observed in the distribution of colocalized puncta along neurites (data not shown). Given the high variability, especially in control cells, we did not detect differences between DMSO and Aβ treatments when focusing on distal sites of Tau-positive neurites either. However, we did observe an accumulation of co-localized events in Aβ-treated cells when neurons were exposed to puromycin for 30 minutes compared to the 5-minute treatment (Fig 5D). We therefore focused on the 30-minute puromycin treatment and analyzed the frequency distribution of translating RNAs, measured as the proportion of colocalized puncta. Interestingly, from all translating RNAs detected, half of them were found within the first 30 μm proximal to the soma in control cells, whereas this proportion was significantly reduced in Aβ-treated cells and consequently the percentage of peripheral translating RNAs increased (Fig 5F and G). Altogether these results show that Aβ oligomers increase the sites of localized translation in distal Tau-positive neurites in line with previously published reports (Baleriola et al., 2014; Li and Gotz, 2017;Walker et al., 2018). Thus, the combination of RNA and protein staining techniques followed by image processing and binarization, and object-based colocalization can be successfully used to detect sites of local RNA translation in neurons which might be important to unravel the extent of local changes in early stages of AD and other neurological diseases.

## Discussion

Highly polarized cells like neurons heavily rely on the asymmetric distribution of their proteome for their functionality. It was classically thought that proteins that support dendritic and axonal functions are synthesized in the soma and then transported to their target destination at peripheral sites of the neuron. This soma-centric view of protein synthesis has slowly changed over the last two decades and it is now accepted that neurites contain mRNAs and components of the translation machinery and are thus able to produce proteins locally. Local protein synthesis enables neurites (both dendrites and axons) to change their proteome in an acute manner in order to adapt to fast environmental changes. Since the first studies that unambiguously demonstrated the existence of local translation in neurons (Koenig, 1967; Giuditta et al., 1968; Steward and Levy, 1982; Torre and Steward, 1992; Feig and Lipton, 1993) most work in the field has focused on understanding the role of locally produced proteins in brain physiology. For example, a subset of mRNAs translated in dendrites, which include CamK2a, Calmodulin or Bassoon, is involved in synaptic plasticity (reviewed in (Holt et al., 2019)). Intra-axonal synthesis of β-actin (Leung et al., 2006), RhoA (Wu et al., 2005) or Par3 (Hengst et al., 2009) is important in growth cone behavior and axon elongation during nervous system development. Proteins involved in mitochondrial function such as LaminB2 (Yoon et al.,2012)or COXIV (Aschrafi et al., 2010) are locally synthesized in axons and contribute to their maintenance in post-developmental stages. However much less is known on the role of local protein synthesis in nervous system pathologies, especially those of the CNS. Alzheimer’s disease (AD), like other neurodegenerative diseases, is characterized by synaptic dysfunction during early stages (Palop and Mucke, 2010). Understanding dynamic early changes in the local proteome (axonal, dendritic or synaptic) is in our view crucial to understand basic pathological mechanisms underlying AD and likely other neurological diseases. Recent work has shown that regulation of intra-axonal protein synthesis induced by Aβ_1-42_ oligomers, whose accumulation is central to AD, contributes to neurodegeneration (Baleriola et al., 2014). These findings support a model in which retrograde transport of locally produced proteins leads to pathological, transcriptional changes in the neuronal soma. More recently, a link between intra-dendritic translation, and Tau mislocalization and hyperphosphorilation has been found (Kobayashi et al.,2017; Li and Gotz, 2017). Thus, dysregulation of the local translatome in neurons might play a more relevant role in AD than previously acknowledged. It is therefore important to know the extent and location of newly-synthesized proteins in order to understand early changes in the AD brain.

Despite local translation is finally being accepted by the scientific community, the accurate measurement of this phenomenon is still challenging partly due to the limited amount of proteins that are locally produced, especially in adult axons (Rangaraju et al.,2017). There are other experimental challenges that will be not be discussed herein since the technicalities are beyond the scope of this manuscript. Our own results show that new-synthesized neuritic proteins measured at distal positions (100-150 μm from the cell nucleus) can be ~20 to 30 times less abundant than those measured in the soma in a 30-minute time frame (Fig 2). Thus, local translation events can be easily overlooked when visualizing *in situ* protein production under the microscope. FUNCAT (Dieterich et al., 2010), and SUnSET (Schmidt et al., 2009) are commonly used techniques in the field of local translation. Both are based on the labeling of newly-synthesized proteins, with noncanonical amino acids in the case of FUNCAT or a tRNA analog in the case of SUnSET. In both cases the noncanonical molecules can be fluorescently tagged. Yet when these methods have proven very helpful to analyze the amount of newly-synthesized dendritic (Dieterich et al., 2010) and axonal (Wong et al., 2017; Walker et al., 2018) proteins measured as fluorescent intensity in labeled cells, discrete foci of newly produced proteins can come unnoticed unless enhanced.

Some variations of the aforementioned techniques such as Puro-PLA or FUNCAT-PLA have been used to accurately measure discrete translation sites of specific proteins along neurites (tom Dieck et al., 2015). Both proximity ligation assays (PLA) are based on the spatial coincidence of two antibodies, one that recognizes the recently synthesized polypeptide chain (anti-puromycin in the case of Puro-PLA; anti-biotin for FUNCAT-PLA) and another one that recognizes a specific protein of interest. However, neither PLA approach is useful to analyze all translation foci. Other modifications of SUnSET have been used to evaluate overall discrete intra-neuritic and intra-dendritic translation events. For example, co-incubation of neurons with both puromycin and the translation inhibitor emetine prior to fixation prevents the puromycilated polypeptide chain release from the ribosomes. This approach is known as ribopuromycilation (RPM) and it allows the visualization of active polyribosomes in the neuronal soma and along neurites (Graber et al., 2013). Additionally, in puromycin-labeled fixed cells, proximity ligation assay (Puro-PLA) employing a single antibody against puromycin has been used to accurately identify discrete local translation sites in dendrites (Rangaraju et al., 2019). Here we describe a strategy to enhance puromycin hotspots in neurites following SUnSET, based solely on image processing and the assisted quantification of the resulting objects. Our technique does not require the incubation of the cells with any translation inhibitor besides puromycin, and it avoids the processing of the samples for proximity ligation assay, which can be pricy and time consuming.

We first performed edge detection to find discontinuities in our puromycin labeling that could result from a punctate staining arising from discrete positive foci. Instead of using the Find Edges command in FIJI/ImageJ which applies a Sobel edge detector, we used the default 5×5 kernel in the convolver which is a Laplacian edge detector instead. Laplacian operators are very accurate in finding edges in an image but also very sensitive to background noise (Bannister and Larkman, 1995a). We therefore adjusted the brightness and contrast of our micrographs after applying the filter in order to eliminate highlighted pixels outside the area established by the neuronal / axonal markers βIII tubulin and Tau. Image processing with the Laplacian operator highlighted events in the periphery of neurons that could be selected and binarized with the MaxEntropy mask (Fig. 3C, 4D, 5E and 5G). Binarization in unprocessed images resulted in the selection of the puromycin signal confined mainly to the cell body region (not shown). To determine if our processing method worked in highlighting local events, we evaluated the effect of Aβ_1-42_ oligomers on hippocampal neurites. Aβ oligomers are known to increase puromycin intensity when applied locally to axons, which reflects changes in local protein synthesis (Walker et al., 2018). In our case, we observed similar results in neurites following bath application of hippocampal neurons with Aβ (Fig 2B): the effect elicited by Aβ was visible beyond the canonical ER domain and did not affect the cell body (Fig 2C and D). These results are compatible with changes in local translation but they do not address whether actual local sites of protein synthesis are affected by Aβ oligomers. Discrete puncta in distal neuritic sites likely reflect foci of localized translation (Graber et al., 2013; Rangaraju et al., 2019). We quantified discrete puromycin-positive foci in distal neuritic sites in response to Aβ_1-42_ with the particle analyzer after image processing with the convolver (assisted quantification). We observed 1) an enhancement of discrete puromycin staining in both DMSO- and Aβ-treated neurites compared to visual inspection of raw puromycin staining (Fig 3B and C), 2) an enhancement of the effect of Aβ on newly-synthesized neuritic proteins compared to controls (Fig 3B, C and H) and 3) a better correlation between the unbiased measurement of puromycin intensity (Fig 2D) and the number of discrete puromycin-positive sites in processed images (Fig 3G). Altogether these results indicate that binarizing images from puromycin-positive cells after applying a Laplacian edge detector allows the assisted quantification of neuritic translation sites yielding results that resemble those obtained from an unbiased measurement of raw puromycin intensity (Fig 2D, 3G and 3H). Additionally, our results unravel a previously unreported effect of Aβ oligomers on discrete translation events in neurites.

Our first approach was performed in βIII tubulin-positive neurites which correspond to both dendrites and axons, but because changes in local neuronal translation upon Aβ treatment were first described in axons (Baleriola et al., 2014), we applied the same processing workflow to neurites stained with the axonal marker Tau. Results were very similar to those obtained for βIII tubulin-positive neurites when cells were fed with puromycin for 30 minutes (Fig 4). Shorter exposures to puromycin were also performed in order to minimize the possible detection of newly-synthesized proteins diffused from the soma. Puromycin pulses as short as 10-15 minutes have been successfully used to detect changes in intra-axonal protein synthesis upon acute exposure of axons to Aβ oligomers (Walker et al., 2018). However, we did not observe changes between DMSO- and Aβ-treated cells possibly due to the slow pace of the translation machinery after a 24 hour treatment.

In light of our results we addressed whether distal puromycin-positive events in neurites arose from localized RNAs to determine if we were actually measuring local protein synthesis. We applied the processing protocol followed for puromycin staining to SYTO-positive neurites. SYTO RNASelect green fluorescent dye selectively binds neuritic RNA (Fig 5A). We observed that Aβ oligomers did not change the distribution of RNA granules along neurites (Fig 5B and E) nor their amount in distal sites (Fig 5C). These results are compatible with other experiments performed in our laboratory aimed at labeling neuritic RNAs with alternative techniques (data not shown). Taking advantage of the fact that SYTO-labelled cells were also labeled with puromycin, after binarizing the images corresponding to both stainings we applied the AND function in the image calculator which essentially retrieves the colocalization between objects. In our system colocalized objects (cyan, Fig 5E) represent sites of actively translating RNAs. We particularly focused on colocalized objects resulting from 30-minute puromycin pulses, which were higher than for shorter puromycin exposures (Fig 5D). Interestingly, Aβ_1-42_ increased the proportion of RNA translation in distal sites of Tau-positive neurites, beyond the ER domain. Thus, the combination of RNA and protein staining techniques followed by image processing and binarization, and object-based colocalization can be successfully used to detect sites of local RNA translation in neurons.

Other edge detectors, Laplacian operators distinct to 5×5 matrices or other background subtraction methods can be used depending on the sample requirements and the researcher’s criteria. However, the image processing approach described herein has proven very useful to detect discrete events with low pixel intensity, which is the expected characteristic of neuritic local translation sites.

Altogether, this study provides a simple method of quantifying local RNA translation sites using object-recognition and object-based colocalization analyses which allows a better understanding the effect of Aβ_1-42_ in neurites.

## Acknowledgments

We thank member of the Neurobiology Lab (Achucarro Basque Center for Neuroscience) for sharing the Aβ peptides with us.

## Author contributions

J.B. conceived the project and designed the experiments. M.G., M.B.U., A.F.R.B., J.I. and J.B. performed experiments. M.G., M.B.U. and J.B. performed data analysis and wrote the manuscript.

